# Disrupting selective persistent activity with electrical stimulation impairs human working memory

**DOI:** 10.1101/2025.08.20.671301

**Authors:** Jonathan Daume, Mar Yebra, Sophia Cheng, Chrystal M. Reed, Ivan Skelin, Yousef Salimpour, Jan Kamiński, Andre Cornejo Marin, William S. Anderson, Taufik A. Valiante, Adam N. Mamelak, Ueli Rutishauser

## Abstract

Working memory (WM) enables the temporary maintenance and manipulation of information, supporting flexible, goal-directed behavior. While converging evidence suggests that persistent activity in the hippocampus and other areas of the brain contributes to WM storage, this link has not been examined by manipulating neural activity experimentally. Here, we combined simultaneous intracranial single-neuron recordings with focal electrical stimulation in the human hippocampus to test the role of memoranda-selective persistent neural activity for WM. Thirty patients with implanted hybrid depth electrodes performed a WM task with images as memoranda. Electrical stimulation (2 s, 50 Hz, 1 mA) was delivered to the hippocampus during the maintenance period on a subset of trials. Behaviorally, stimulation impaired WM performance, increasing response times especially in low-load conditions. Neuronally, stimulation reduced memoranda-selective activity in hippocampus and ventral temporal cortex (VTC), thereby disrupting content-specific neural representations of WM content. The extent of stimulation-induced change in firing rate mediated trial-by-trial impaired WM-related behavior, linking disrupted neural selectivity to impaired WM. At the population level, stimulation shifted neural trajectories farther from attractor states, consistent with degraded mnemonic fidelity. Together, these data provide evidence that selective persistent activity of individual neurons in hippocampus and VTC supports WM maintenance in humans. Our results demonstrate that hippocampal stimulation disrupts both single-neuron coding and population-level attractor stability, linking cellular mechanisms to behavior and highlighting the contributions of persistent activity to WM maintenance.

## Introduction

Working memory (WM) is a fundamental cognitive process that enables the temporary storage and manipulation of information relevant to current goals (Baddeley and Hitch 1974; Cowan 2010; Baddeley et al. 2020). In humans, neurons in the human hippocampus and other medial temporal lobe areas such as the amygdala exhibit WM-content selective persistent activity during the WM maintenance period, a hallmark neural signature thought to reflect the online storage of task-relevant information in WM (Kamiński et al. 2017, 2020; Kornblith et al. 2017; Boran et al. 2019; Daume et al. 2024b, a; Paluch et al. 2025). Such neurons show elevated and sustained firing rates that are selective for the stimulus held in memory, persisting in the absence of sensory input. Memoranda-selective persistent firing is thought to be a key neural mechanism underlying WM (Curtis and D’Esposito 2003; Sreenivasan et al. 2014; Constantinidis et al. 2018; Kamiński and Rutishauser 2020; Curtis and Sprague 2021; Wang 2021). However, to date, evidence for a role of selective persistent activity in the hippocampus and other areas in the brain in WM remains correlational, with no tests of this relationship using perturbation methods done so far.

Here, we address this question by leveraging a rare opportunity to conduct simultaneous intracranial single-neuron recordings in multiple brain areas and focal electrical stimulation in the hippocampus in human patients undergoing invasive monitoring for epilepsy. This combination of techniques allowed us to test whether perturbing neural activity during the maintenance phase of a WM task would modulate either neural representations of mnemonic content in various cortical and subcortical areas of the brain and/or behavioral performance. This approach allows us to directly interfere with a candidate mechanism of information maintenance (selective persistent activity) and to measure the downstream neural and/or behavioral effects of this manipulation. WM maintenance through persistent activity is thought to be supported by a distributed network of brain areas rather than a single brain area, a theory supported by the widespread observation of selective persistent activity in different brain areas across species that include the hippocampus, amygdala, posterior temporal cortex, dorsolateral pre-frontal cortex in the frontal lobe, and, in mice, anterior lateral motor cortex (Fuster and Jervey 1982; Miller et al. 1991; Constantinidis et al. 2001; Kamiński et al. 2017; Inagaki et al. 2019; Daume et al. 2024b, a). As such, we would expect stimulation of a single node in this network to disrupt persistent activity in several brain areas. Here, we test this theory by stimulating the hippocampus and assessing selective persistent activity at the single neuron level in several temporal and frontal brain areas.

In parallel to single-neuron analyses, we also examined the structure of neural population activity in low-dimensional state space. Prior work has suggested that during WM, neural population dynamics evolve toward and remain near fixed point attractor states, i.e., stable configurations that correspond to specific contents held in memory (Wang 2001; Wimmer et al. 2014; Panichello et al. 2019; Kamiński and Rutishauser 2020; Penny 2024). The stability of these attractors has been linked to WM performance: the closer the population trajectory remains to the attractor state, the more accurate the behavioral response (Kamiński et al. 2017). Disrupting selective persistent activity during WM maintenance, therefore, would be predicted not only to affect individual neurons, but also to alter the geometry of population-level dynamics, pushing neural trajectories away from their optimal attractor states.

Based on this theoretical framework, we tested two main hypotheses. First, if selective persistent activity in medial temporal lobe as well as posterior-temporal and frontal cortical areas supports WM maintenance, then disrupting that activity with electrical stimulation should impair WM-dependent behavior and reduce stimulus-specific firing in individual neurons. Second, if population activity supports WM through attractor dynamics, then stimulation should distort these trajectories and increase the distance from the appropriate attractor state, reflecting degradation of mnemonic fidelity.

To test these hypotheses, we asked human epilepsy patients to perform a modified Sternberg task involving WM for visual stimuli, during which we recorded spiking activity from implanted microelectrodes. This task is identical to that used in our prior studies (Daume et al. 2024b, a), in which we demonstrated robust persistent activity by visually selective ‘category’ neurons, the activity of which we will analyze here. On a random subset of trials, electrical stimulation was delivered focally directly to either the left or the right (but not both) hippocampus during the WM maintenance period. Patients did not know in which trials stimulation was applied, allowing for within-subject comparisons of neural activity and behavior between stimulated and non-stimulated conditions. This design enabled us to ask: (1) whether electrical stimulation, and if so for what kind of trials, modulates behavioral performance when applied during the WM maintenance period, (2) whether electrical stimulation affects category-selective firing in single neurons and if so in which brain area(s), and (3) whether such stimulation alters the structure of population-level dynamics in a way consistent with disruption of attractor states. To our knowledge, this is the first study to combine electrical stimulation and simultaneous single-unit recordings to test the role of selective persistent activity in human WM.

## Results

### Task, behavioral results, and electrophysiology

30 patients (32 sessions; **Table S1**) performed the task. Stimulus material was pictures belonging to one of five different categories (people, animals, cars/tools (depending on variant), fruit, and landscapes; see also (Daume et al. 2024b, a)). In each trial, subjects were first shown either one (load 1) or three (load 3) consecutively presented pictures. Following a maintenance period of 2.5–2.8s length, subjects then indicated whether a probe stimulus shown was identical to one of the item(s) they just saw or not (**Fig. 1a**). In a random subset of trials, we delivered electrical stimulation pulses (2 s, 100 bipolar pulses at 50 Hz, see Methods) to the two most medial macro electrodes of an electrode implanted in the left or right hippocampus (**Fig. 1c**; **Table S1;** see Methods) during the maintenance period (**Fig. 1a**, arrow). Patients did not know in which trials they received the stimulation and did not report any effects of the stimulation.

**Figure 1.**
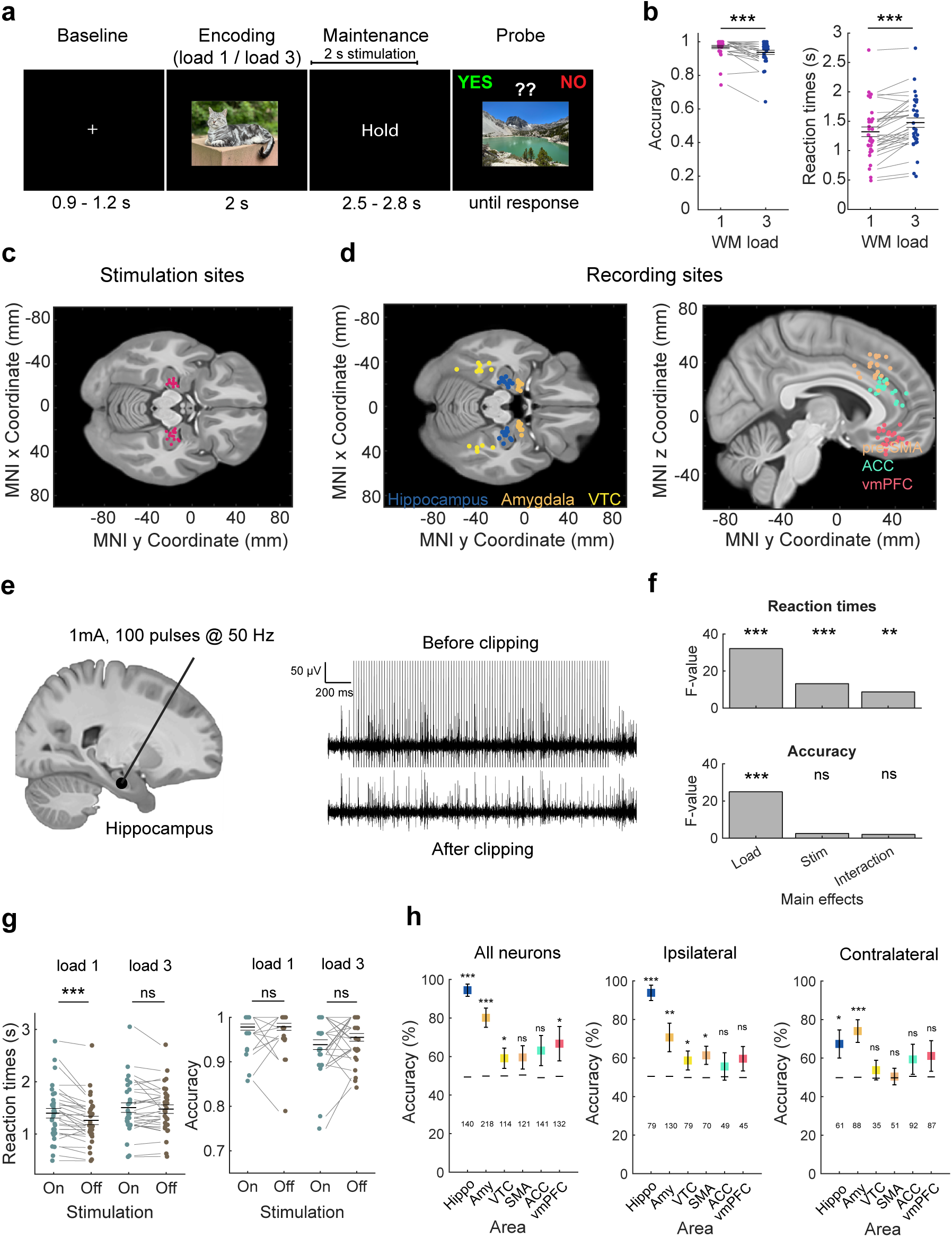
Task, recordings sites, stimulation properties, behavior, and population decoding. **(a)** Task. Patients performed a modified Sternberg WM task, encoding either one (load 1) or three (load 3) novel images drawn from five visual categories. Following a variable delay (maintenance, 2.5-2.8s), a probe image was presented, and participants indicated whether it matched one of the encoded images or not. Electrical stimulation (2 s, 50 Hz) was applied in a subset of trials during the WM maintenance period. **(b)** Behavioral performance. Participants showed higher accuracy (left) and faster RTs (right) in load 1 compared to load 3 trials (all trials). **(c)** Subset of hippocampal electrodes that were stimulated across all participants. **(d)** Recording locations across six brain regions: hippocampus, amygdala, VTC, pre-SMA, ACC, and vmPFC. Each dot represents one microwire bundle. Coordinates are plotted in MNI152 space. **(e)** Electrical stimulation was applied via the two most medial macro contacts of the hybrid electrode implanted in either the left or right hippocampus (bipolar, 1 mA, 100 biphasic pulses at 50 Hz over 2 s). Example trace in the lower panel shows removal of stimulation artifacts using interpolation from –0.5 to +2.0 ms around each pulse. This preprocessing step enabled analysis of neural activity during stimulation. Artifact removal was performed in all brain regions (see Methods). The same interpolation windows and removal processes were applied to non-stimulation trials to ensure unbiased comparisons. **(f,g)** Behavioral effects of stimulation. **(f)** For reaction times, we observed a significant main effect for *load* and *stimulation* as well as a significant interaction (top). Accuracies only showed a significant main effect for *load* but no significant effect for *stimulation* and no significant interaction. **(g)** Hippocampal stimulation impaired WM performance as measured by RTs specifically in load 1 trials, resulting in slower RTs in *stimOn* versus *stimOff* trials (left). Reaction times in load 3 trials or accuracies (right) in either load condition did not differ significantly between stimulation conditions. **(h)** Decoding of stimulation condition (*stimOn* vs. *stimOff*) from population activity. A linear SVM decoder was trained to classify stimulation condition based on FRs during WM maintenance. Decoding was performed separately for all neurons (left), ipsilateral neurons (middle), and contralateral neurons (right) across regions. Significant decoding was strongest in hippocampus and adjacent areas. Squares indicate mean decoding accuracy; error bars reflect standard deviation across 50 subsamples. Dashed lines indicate the mean across 500 shuffled permutations. Numbers below bars indicate neuron counts. *** p < 0.001, ** p < 0.01, * p < 0.05, ns = not significant, permutation-based t-test.

Patients performed well, with a mean accuracy of 95.32 ± 6.17 % (mean ± standard deviation (SD)). Across all sessions, subjects responded slower (1.48 s vs 1.32 s; t(31) = 5.97; p < 0.001) and less accurate (93.73 % vs 96.89 %; t(31) = -4.95; p < 0.001) in load 3 as compared to load 1 trials (**Fig. 1b**).

We next tested whether stimulation of the hippocampus during the WM maintenance affected behavioral performance, which we assessed with both accuracy and reaction time (RT). We used a permutation-based 2×2 analysis of variance (ANOVA) analysis with factors *load* (load 1 vs load 3) and *stimulation* (stimOn vs stimOff) and RT as the dependent variable. This analysis revealed significant main effects for both *load* (**Fig. 1f top**; F(1,30) = 32.08; p < 0.001) and *stimulation* (F(1,30) = 13.20; p < 0.001) and a significant interaction (F(1,30) = 8.73; p = 0.005; one patient with an accuracy of less than 65% in stimulation trials was excluded from analyses of stimulation effects, but including this patient did not change the result). Post-hoc permutation-based t-tests revealed that RTs were significantly slower in *stimOn* trials compared to *stimOff* trials in load 1 (**Fig. 1g left**; t(30) = 4.48, p < 0.001) trials, with no significant difference in load 3 trials (t(30) = 1.04, p = 0.32). In contrast, repeating the same 2×2 ANOVA for accuracy revealed only a significant main effect for *load* (**Fig. 1f bottom**; F(1,30) = 24.93; p < 0.001) but not for *stimulation* (F(1,30) = 2.55; p = 0.12; **Fig. 1g right** shows the effect of stimulation accuracy in each load condition), with no significant interaction (F(1,30) = 2.06; p = 0.16). This analysis of the behavior therefore revealed that electrical stimulation of the hippocampus disrupted WM-related behavior specifically in load 1 trials, in which stimulation led to slower RTs but did not affect accuracy.

Subjects with epilepsy have different degrees of hippocampal sclerosis. We next asked whether the effect of hippocampal stimulation we found in load 1 trials was different as a function of the extent of sclerosis of the stimulated hippocampus (see methods and **Table S1**). This was not the case: A mixed-effects GLM with factors sclerosis rating and stimulation on/off revealed no significant interaction between stimulation and sclerosis rating (**Fig. S1a**). In addition, we tested whether the anatomical location of stimulation along the x-, y-, or z-axis in MNI space modulated the magnitude of the RT effect but found no evidence for such an association (**Fig. S1b**).

While patients performed the task, we recorded single neuron activity in six different brain regions: hippocampus, amygdala, ventral temporal cortex (VTC) (Wadia et al. 2026), pre-supplementary motor area (pre-SMA), dorsal anterior cingulate cortex (dACC), and ventromedial pre-frontal cortex (vmPFC; **Fig. 1d**). We acquired high-quality simultaneous single neuron recordings during stimulation in a subset of n = 15 sessions (14 patients) due to technical limitations (see Methods). Across the subset of sessions with both stimulation and recordings, we recorded in total 866 putative single neurons (140 in the hippocampus, 218 in the amygdala, 114 in VTC, 141 in dACC, 121 in pre-SMA, and 132 in vmPFC.

To assess the overall impact of electrical stimulation of the hippocampus on neural activity in each recorded brain area, we employed a linear decoder to classify *stimOn* versus *stimOff* trials based on the firing rates of all neurons within a given brain region during the maintenance period. As expected, stimulation exerted the most pronounced effects in the hippocampus and the adjacent amygdala (**Fig. 1h**). Additionally, the decoder was able to differentiate between *stimOn* versus *stimOff* trials from neural activity in both the VTC and vmPFC across both hemispheres. Notably, robust decoding in both the ipsilateral and contralateral hemispheres relative to the stimulation site was observed only in the hippocampus and amygdala (**Fig. 1h**). Note that the purpose of this analysis was solely to identify brain regions in which hippocampal electrical stimulation produced detectable changes in neural activity, irrespective of the specific nature of those changes. The results of this analysis do not form the basis for subsequent investigations, which focus exclusively on the subset of neurons that are category selective.

### Category neurons maintain mnemonic information through selective persistent activity

Before examining how electrical stimulation affects the activity of single neurons, we first identified where in the brain neurons encoded the visual category of presented stimuli and whether these neurons remained persistently active during WM maintenance. We focused our analysis on neurons whose firing rates following stimulus onset varied significantly as a function of the visual category of the image shown on the screen. Throughout the manuscript, we refer to these as *category neurons*.

To identify category neurons, we analyzed spike counts within a 200–1200 ms window following picture onset (during encoding phases 1–3 and the probe phase). We applied a one-way ANOVA to test for significant modulation by stimulus category (five levels), followed by a right-tailed, permutation-based t-test comparing the neuron’s response to its most preferred category (i.e., the category eliciting the highest average spike count) against its responses to all other categories. Neurons that passed both statistical tests (p < 0.05 for both tests) were classified as category neurons, with the preferred category defined as the category to which their response is maximal (see Methods for details). Significant proportions of category neurons were found across all recorded regions, with the highest prevalence observed in the hippocampus, amygdala, VTC, and vmPFC (**Fig. 2a**; example neurons shown in **Fig. 2c–f**).

**Figure 2.**
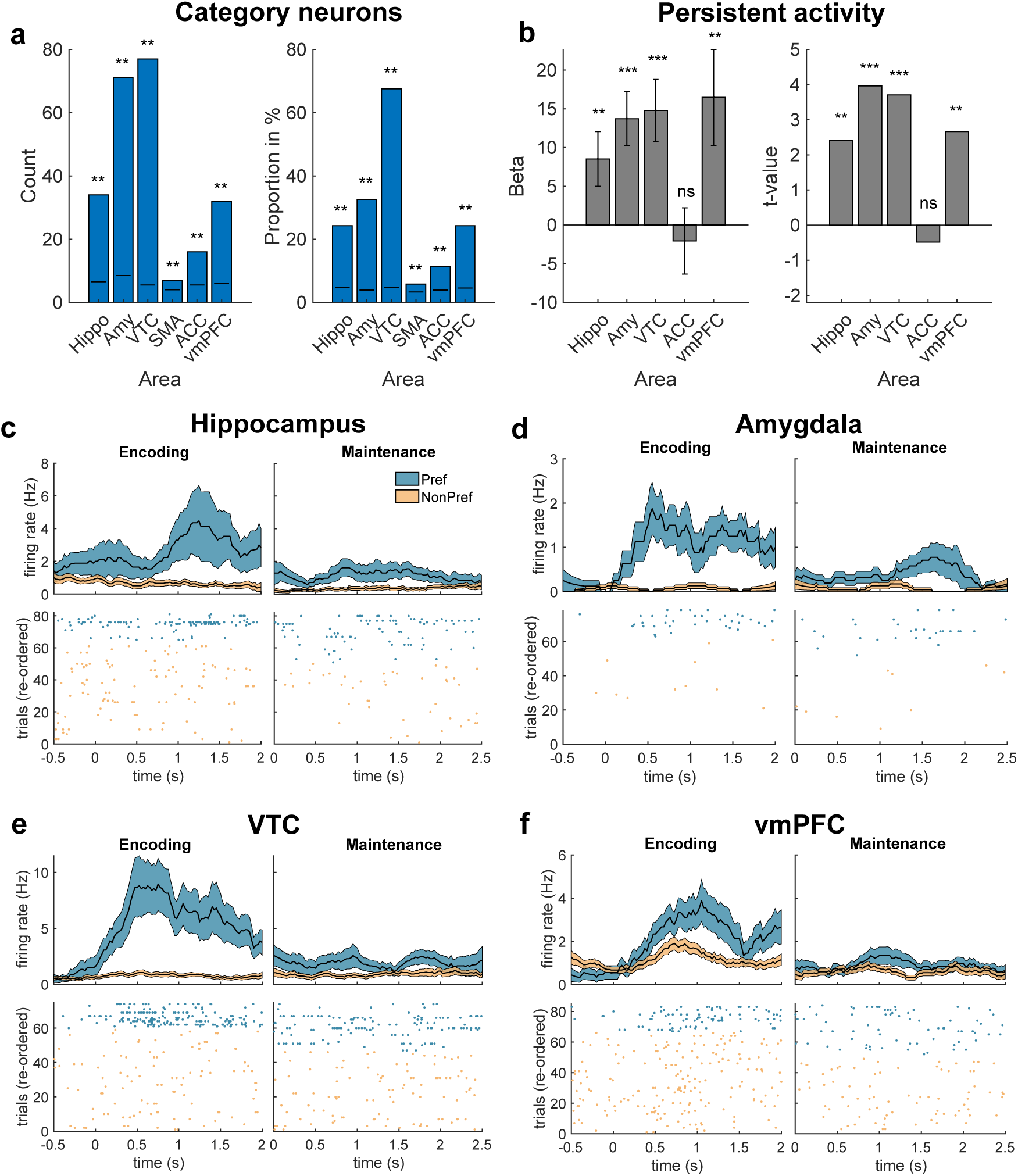
Category selectivity and persistent activity during WM maintenance. **(a)** Count (left) and proportion (right) of category-selective neurons (“category neurons”) identified in each recorded brain area during picture presentation. Category neurons were most prevalent in hippocampus, amygdala, VTC, and vmPFC. Black lines within each bar represent the 99th percentile of a null distribution obtained by randomly shuffling category labels across 500 iterations. **(b)** Persistent activity of category neurons during the WM maintenance period in non-stimulated trials. Mixed-effects GLMs with neuronID nested into patientID (see Methods) revealed significantly higher firing rates when images of the neurons’ preferred categories were held in WM, with the strongest effects observed in hippocampus, amygdala, VTC, and vmPFC. Pre-SMA was excluded from this analysis due to an insufficient number of identified category neurons for reliable statistical inference. **(c–f)** Example category neurons from (c) hippocampus, (d) amygdala, (e) VTC, and (f) vmPFC. Shown are average firing rates (top) and trial-by-trial raster plots (bottom) during encoding and maintenance, separated by preferred (blue) and non-preferred (orange) category conditions. Only stim off trials are shown. *** p < 0.001, ** p < 0.01, * p < 0.05; ns = not significant.

To determine whether identified category neurons exhibited persistent activity during the WM maintenance period (which is statistically independent; neurons were selected during the encoding period), we fit mixed-effects general linear models (GLMs) with *neuronID* nested into *patientID* separately for each brain region (see Methods for details). The models included category preference (two levels: preferred vs. non-preferred category) as a predictor of baseline-corrected FRs during the maintenance period, considering only correct *stimOff* trials. We excluded the pre-SMA from this analysis due to the low number of identified category neurons in this region (<10), which precluded reliable statistical inference.

Across all examined regions except the dACC, we found significantly higher firing rates during maintenance when neurons’ preferred categories were held in WM compared to non-preferred categories **(Fig. 2b**; p < 0.01 for all regions). These results support the role of category neurons in maintaining category-specific information during the maintenance period of the task, independently confirming this finding in an entirely new dataset (Daume et al. 2024b, a). Apart from confirming earlier findings, here we demonstrate, for the first time, persistent activity of category**-**selective neurons in visual (VTC) (Wadia et al. 2026) and prefrontal (vmPFC) (Aquino et al. 2023) areas of the human brain.

### Electrical stimulation disrupts selective activity in hippocampus and VTC

Given that electrical stimulation of the hippocampus impaired WM behaviorally (see **Fig. 1f,g**), we hypothesized that stimulation disrupted the neural mechanisms thought to underlie information storage in WM: selective persistent activity of category neurons. To test this hypothesis, we conducted nested mixed-effects GLMs separately for each brain region (excluding pre-SMA due to insufficient neuron count, see above). The models included *preference* (preferred vs. non-preferred), *stimulation* (stimOn vs. stimOff), and their interaction. The dependent variable was the baseline-corrected FR of category neurons during the WM maintenance period. We predicted that stimulation would disrupt persistent activity, evidenced by a significant interaction between *preference* and *stimulation*.

This analysis revealed a significant interaction between *preference* and *stimulation* in two brain areas: the hippocampus and VTC (**Fig. 3a**), but not the amygdala, vmPFC, and dACC (**Fig. S2**). To further examine this effect, we next focus only on the neurons in the hippocampus and VTC, where stimulation had a significant effect. We compared FRs during *preferred* versus *non-preferred trials separately for stimOn* and *stimOff* trials at the single-neuron level, combining neurons from the hippocampus and VTC. During *stimOff* trials, as expected, neurons exhibited significantly higher FRs in preferred compared to non-preferred trials (**Fig. 3b right**; beta = 12.16; t = 4.34; p < 0.001; nested mixed-effects GLM). In contrast, during *stimOn* trials, category selectivity was abolished, with no significant difference between *preferred* and *non-preferred* trials (**Fig. 3b left**; beta = 1.11; t = 0.35; p = 0.72), indicating a loss of selective persistent activity in *stimOff* trials.

**Figure 3.**
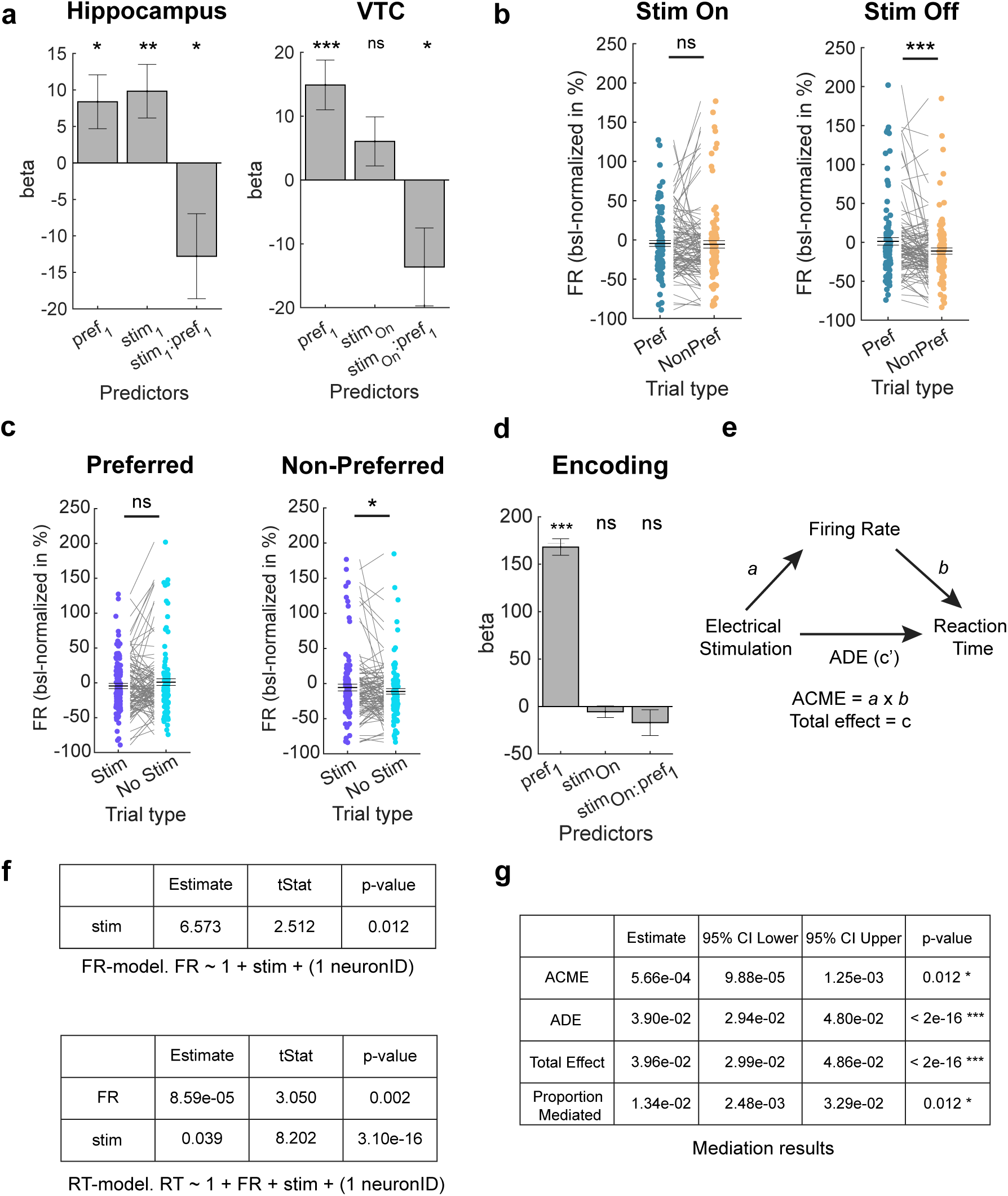
Electrical stimulation disrupts persistent activity of category neurons. **(a)** Mixed-effects GLM results in hippocampus (left) and VTC (right) revealed a significant interaction between preference and stimulation, indicating that persistent category-selective activity was disrupted by hippocampal stimulation across both load conditions. See **Fig. S2** for results from other areas. **(b, c)** Post-hoc comparisons of baseline-normalized FRs of category-selective neurons pooled across hippocampus and VTC. Statistical comparisons were performed using mixed-effects GLMs. **(b)** During stimOff trials, FRs were significantly higher when the preferred category was maintained in WM compared to non-preferred trials, reflecting category-selective persistent activity. This difference was abolished during stimOn trials. **(c)** Direct comparisons between stimulation conditions revealed that FRs were significantly increased in stimOn relative to stimOff trials specifically for non-preferred trials, whereas no significant difference was observed for preferred trials. **(d)** GLM analysis during the encoding period (pre-stimulation) showed no significant interaction between stimulation and preference, confirming that observed effects were specific to the maintenance window. **(e)** Schematic of the mediation framework. This analysis tests whether stimulation-induced changes in FR of category-selective neurons mediate the effect of hippocampal stimulation on RT. **(f)** Structure of the mediation analysis. Two GLMs were used: an FR-model quantifying the effect of stimulation on FR (mediator), and an RT-model quantifying the joint effects of stimulation and FR on RT (outcome). **(g)** Results of the mediation analysis. ACME was significant, indicating that stimulation-induced changes in FR significantly mediated the effect of stimulation on RT in non-preferred trials of the load 1 condition. The average direct effect (ADE) remained significant, suggesting that additional factors also contribute to the behavioral effect. In **(b, c**), data points represent single-neuron averages across trials for visualization only. Statistical analyses were performed using mixed-effects GLMs with neuronID nested within patientID. **\***\*\*p < 0.001, **p < 0.01, * p < 0.05; ns = not significant; mixed-effects GLMs and permutation-based t-test.

What caused the loss of category selectivity during stimulated trials? This effect was primarily driven by a selective increase in firing rates in *stimOn* as compared to *stimOff* trials for *non-preferred* trials (**Fig. 3c right;** beta = 5.46; t = 2.30; p < 0.05), whereas firing rates did not differ significantly between *stimOn* and *stimOff* trials for *preferred* trials (**Fig. 3c left;** beta = -5.38; t = -1.41; p = 0.16). Separating trials into the load 1 and 3 condition, this effect was only significant in load 1 trials (stimOn vs stimOff, preferred: beta = 1.74; t = 0.28; p = 0.78; non-preferred: beta = 6.82; t = 2.37; p < 0.05) but not load 3 (*stimOn* vs. *stimOff*, *preferred*: beta = -7.48; t = -1.60; p = 0.11; *non-preferred*: beta = 2.97; t = 0.70; p = 0.49). Together, this analysis shows that category selectivity was lost because stimulation increased firing rates during non-preferred trials.

This interaction was specific to the maintenance period and was not observed during the encoding phase, i.e., when no stimulation occurred (GLM interaction p = 0.21; **Fig. 3d**). Additionally, including *stimulation site* (ipsilateral vs. contralateral) as a factor in the model did not yield a significant three-way interaction (*preference* × *stimulation* × *site*: p = 0.85), suggesting that the disruption of persistent activity was comparable for neurons recorded ipsi- and contralateral to the stimulation site.

### Firing rates mediate behavioral effect of hippocampal stimulation

So far we showed that electrical stimulation of the hippocampus caused both a neural and a behavioral effect: it disrupted selective persistent activity of category-selective neurons in the hippocampus and VTC, and it impaired WM-related behavior in load 1 trials (**Fig. 1g**). Where these two effects related to each other? To answer this question, we next asked whether the behavioral effects of stimulation were mediated by stimulation-induced changes in the FR of category-selective neurons. Demonstrating such mediation would support a direct link between changes in neuronal firing and behavior. To this end, we performed a mediation analysis (MacKinnon et al. 2007; Imai et al. 2010; Tingley et al. 2014) in which *stimulation* served as the treatment variable, *FR* as the mediator, and *RT* as the behavioral outcome (see schema in **Fig. 3e**). Because stimulation-related behavioral effects were only significant in load 1 trials, and stimulation-related changes in FR were confined to non-preferred trials of category cells, we used the non-preferred trials from the load 1 condition for this analysis (see Methods for details).

The mediation framework consisted of two GLMs: one modeling the effect of stimulation on FR (“FR-model), and a second modeling the joint effects of stimulation and FR on RT (“RT-model”). The results of these models are shown in **Fig. 3f**. Stimulation exerted a significant positive effect on FR (**Fig. 3f, top**), indicating that hippocampal stimulation increased neuronal firing during the WM maintenance period. In addition, both stimulation and FR had significant positive main effects on RT (**Fig. 3f, bottom**), indicating that stimulation prolonged RTs and higher FRs were associated with longer RTs on a trial-by-trial basis.

Combining these models in the mediation analysis revealed that stimulation had a robust direct effect on RT, reflected in a significant average direct effect (ADE; **Fig. 3g**). FR of category-selective neurons in the hippocampus and VTC in addition also significantly mediated the stimulation-induced change in RT, as indicated by a positive average causal mediation effect (ACME; **Fig. 3g**). Thus, a portion of the behavioral impact of hippocampal stimulation was transmitted indirectly through its effect on neuronal firing rates. As expected, the indirect effect was smaller than the direct effect, likely reflecting the limited subset of neuronal activity included in the analysis (see Discussion).

To assess whether the observed indirect effect was specific to the FR of category-selective neurons, we repeated the mediation analysis using the FR of all non-selective neurons in the hippocampus and VTC as the mediator. This analysis revealed no significant ACME for this neuronal population (**Fig. S2d**), indicating that the indirect effect of hippocampal stimulation on RT was specific to category-selective neurons.

We emphasize that these results do not imply that changes in the FR of category-selective neurons fully account for stimulation-induced alterations in WM-related behavior. Rather, they demonstrate that FR contributes directly, at least in part, to the observed behavioral effects. Other neural mechanisms, including changes in temporal firing patterns or LFP-related activity, may also play a role and remain important targets for future investigation.

### Hippocampal stimulation disrupts attractors in state space during WM maintenance

We next investigated how electrical stimulation of the hippocampus disrupted population-level representations (considering all recorded neurons from hippocampus and VTC) of WM content in neural state space. Prior work has demonstrated that during WM maintenance, the neural trajectory of population activity converges near a stable fixed point in state space corresponding to the information held in memory—commonly referred to as an attractor state (Wang 2001). The stability of this trajectory, particularly its proximity to the attractor, has been associated with WM performance: the closer the trajectory remains to the attractor center, the better the performance (Kamiński et al. 2017). Based on our behavioral findings that stimulation impairs WM (manifested as increased RTs and reduced accuracies) we hypothesized that electrical stimulation results in neural population trajectories to deviate from their optimal attractor states. Given that stimulation effects on persistent activity were specific to hippocampus and VTC, we focused this analysis on neurons recorded from those regions.

To examine neural dynamics in state space, we applied demixed principal component analysis (dPCA) as a dimensionality reduction technique (Kobak et al. 2016), projecting trial-wise neural activity into a lower-dimensional space. To avoid ambiguity from multi-item interference, this analysis was restricted to correct load 1 trials only. Importantly, the projection matrix was derived solely from neural activity during the encoding period, ensuring that trajectories during the maintenance period remained independent for subsequent testing. dPCA was applied with picture category as the marginalized variable.

Consistent with prior findings (Kamiński et al. 2017), we identified a four-dimensional subspace (defined by components 1, 2, 3, and 5) that accounted for the highest proportion of variance attributed to category identity (**Fig. 4a**), collectively explaining 60.26% of category-related variance. Neural trajectories corresponding to different categories were clearly separated in this space (**Fig. 4b–d**; in **c**,**d**, only 4 of the 5 categories are shown to increase visibility), and category identity could be significantly decoded from all time points in the first component alone (**Fig. 4b**; see **Fig. S3** for other areas). To assess whether stimulation disrupted proximity to attractor states, we quantified the distance of each trial’s neural trajectory from the relevant attractor. Attractor locations were defined using the mean activity during the maintenance period of *stimOff* trials, where non-interrupted neural trajectories should reside in an optimal state. For each trial, we calculated the distance-to-attractor (DA) as the Euclidean distance to its corresponding category attractor, normalized by the mean distance to all other attractors (Kamiński et al. 2017) (see Methods). A DA value of 1 indicates equidistance to the correct vs. all other attractors, and a DA value of <1 indicates that the neural state is closest to the correct attractor state. Comparing DA values between *stimOn* and *stimOff* trials revealed that electrical stimulation significantly increased the distance of neural trajectories from the attractor (**Fig. 4e**; t(42.01) = 4.10, p < 0.001, unpaired permutation test), indicating a disruption of population-level coding during WM maintenance.

**Figure 4.**
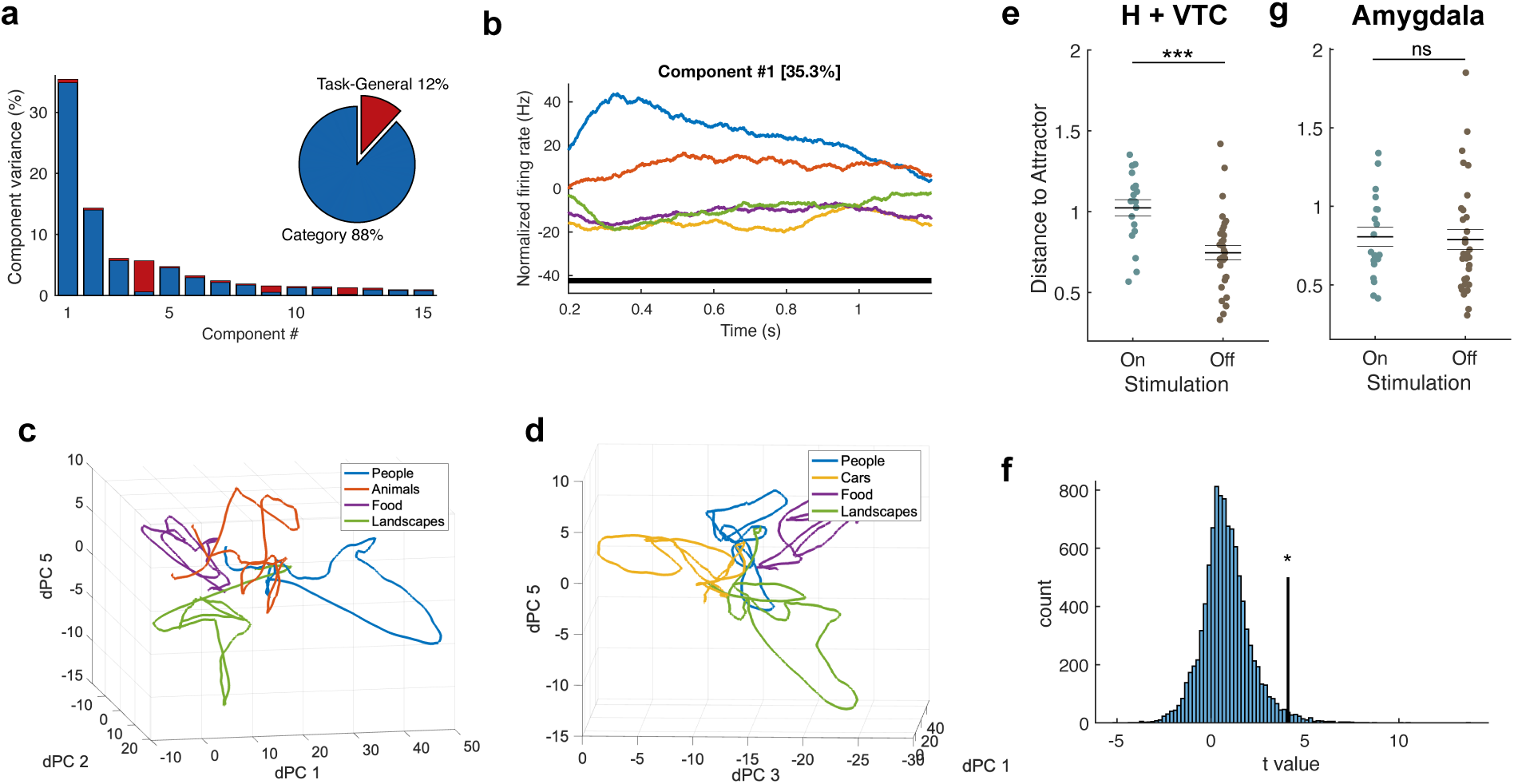
Electrical stimulation disrupts population-level WM coding in state space. **(a)** Variance **in neuronal activity from all recorded neurons in hippocampus and VTC** explained by each dPCA component during encoding. **(b)** Average firing rates projected onto the first dPC, showing clear separation by category (each color is a different category). Black dots at the bottom indicate significant decoding of category at each time point. t = 0 indicates picture onset. **(c–d)** Low-dimensional neural trajectories for different visual categories during encoding (same time window as in panel b), projected into dPC state space. Left and right panels show different combinations of dPC components and categories for better visualization. **(e)** Distance-to-attractor (DA) was significantly higher in *stimOn* versus *stimOff* trials for hippocampus and VTC, indicating disruption of attractor dynamics. **(f)** Permutation test validating that the observed DA shift exceeded what would be expected by chance. **(g)** No significant difference in DA was observed in the amygdala, confirming the regional specificity of the stimulation effect. ***p < 0.001, * p < 0.05; ns = not significant; permutation-based t-test.

A potential concern in this analysis is the introduction of selection bias due to defining attractor states based exclusively on *stimOff* trials, which could artificially favor smaller DA values for those same trials. Thus, to assess whether our findings were driven by this potential bias, we performed a control analysis in which attractor states were redefined 10,000 times using randomly sampled trials drawn from the full dataset, preserving the same distribution of trial counts as in the original analysis. We then recomputed the DA difference between the trial group used to define the attractor (i.e., analogous to the *stimOff* trials in the original analysis) and the comparison group (analogous to the *stimOn* trials) in each iteration. The effect observed using the original *stimOff*-based attractor was significantly larger than the the DA difference obtained by the random permutations (**Fig. 4f**; p = 0.019), demonstrating that the original result was unlikely to be explained by biased attractor selection.

To further validate the specificity of this effect, we repeated the same analysis in all other recorded brain regions. Of these, only the amygdala exhibited robust above-chance category decoding in the first dPCA component (**Fig. S3**). However, unlike in hippocampus and VTC, DA values in the amygdala did not differ significantly between *stimOn* and *stimOff* trials (**Fig. 4g**, t(42.29) = -0.32, p = 0.76). This regional specificity reinforces the conclusion that electrical stimulation selectively disrupts attractor dynamics in hippocampus and VTC, and further rules out the possibility that the observed effects are artifacts of how attractor states were defined. These findings suggest that electrical stimulation of the hippocampus selectively disrupts the fidelity of population-level WM representations in hippocampus and VTC by shifting trajectories away from their category-specific attractor states.

## Discussion

In this study we move beyond correlative observations by combining simultaneous intracranial single-neuron recordings with focal electrical stimulation. Our main contribution is to show that disrupting selective persistent activity with electrical stimulation impairs the maintenance of information in WM. Our results demonstrate that hippocampal stimulation during the WM maintenance period leads to (1) degraded behavioral performance, (2) elimination of selective persistent activity in category neurons in hippocampus and VTC, and (3) distortions in the geometry of neural population dynamics, specifically an increased distance from category-specific attractor states in neural state space. These findings contribute novel insights into the role of the hippocampus and VTC in WM, reinforcing the notion that these areas contribute to maintaining short-term mnemonic representations. Prior studies have documented persistent hippocampal activity during WM tasks and linked it to trial-by-trial performance (Kamiński et al. 2017; Kornblith et al. 2017; Daume et al. 2024b, a), yet such work remained inherently correlational. Lesion studies and observational electrophysiology provide additional evidence for a hippocampal involvement in WM but do not establish a direct link between selective persistent activity and WM specifically (Jeneson and Squire 2012; Goodrich et al. 2019; Borders et al. 2021; Xie et al. 2023; Yonelinas et al. 2024). Similarly, while local pharmacological manipulations can impair persistent activity in animal models, these manipulations are long-acting and do not typically result in behavioral impairment (Wang et al. 2013; Major et al. 2015). Systemic pharmacological manipulations, on the other hand, affect both memory-related behaviors and neural activity, but their systematic and long-acting nature make it challenging to specifically related the observed behavioral impairment to specific aspects of neural activity (Zhou et al. 2011; Vijayraghavan and Everling 2021; Gedankien et al. 2023). Our study fills this gap by showing that brief electrical stimulation of the hippocampus only during the memory period impairs both the neural correlates of WM and behavioral accuracy, including showing that the more selective activity is disrupted, the more behavior is impaired.

### Role of category selectivity during working memory maintenance

Our results demonstrate that hippocampal stimulation alters the neural representation of WM content by reducing category selectivity of single neurons. In the absence of stimulation, category-selective neurons in the hippocampus and VTC exhibited higher firing rates when stimuli from their preferred category were maintained in WM compared to non-preferred categories, consistent with prior work linking such differential activity to WM content (Daume et al. 2024b, a). During stimulation, however, this distinction between preferred and non-preferred conditions was markedly reduced.

Surprisingly, this reduction in selectivity was not driven by a suppression of firing during preferred trials. Instead, stimulation predominantly increased firing rates during non-preferred trials, while firing during preferred trials was only weakly affected. As a consequence, the difference in firing rates between preferred and non-preferred conditions was attenuated, effectively degrading the WM-content specificity of the neural signal during WM maintenance. Thus, stimulation did not abolish persistent firing itself but reduced the contrast between preferred and non-preferred neural states that normally conveys category-specific WM information. We note that similar effects of manipulations on the activity during non-preferred trials have also been seen with pharmacological manipulations of persistent activity: some drugs primarily affecting activity for the preferred stimuli and others for the non-preferred stimuli of persistently activity neurons (Gamo et al. 2010; Major et al. 2018), with both modulating memory as a function of the strength of the difference between the two.

One possible explanation for this asymmetry is that category-selective neurons may be less susceptible to stimulation when they are already actively engaged in representing WM content. During preferred trials, these neurons exhibit elevated and sustained firing, potentially reflecting a stabilized network state that is more resistant to external perturbation. In contrast, during non-preferred trials, i.e., when neurons are closer to baseline firing levels and not actively contributing to WM maintenance, the sort of weak electrical stimulation we used may more readily induce increases in firing. This interpretation suggests that the impact of stimulation depends on the functional state of the neuron at the time of perturbation. This account remains theoretical and awaits experimental testing to determine whether task engagement indeed confers resilience against (weak) electrical stimulation.

We propose that the stimulation-induced increase in firing during non-preferred trials leads to a degradation of the neural code supporting WM content. Under normal conditions, category-selective neurons signal the presence of a preferred stimulus in WM through elevated firing relative to baseline. By elevating firing rates in non-preferred conditions, stimulation may cause these neurons to emit activity patterns that are more ambiguous and less informative with respect to the maintained category. This reduction in the separability of preferred and non-preferred neural states could impair downstream readout mechanisms, thereby increasing uncertainty about whether a probe stimulus matches the maintained item. Such degraded neural representations provide a plausible account for the observed prolongation of RTs in stimulated trials, particularly in low-load conditions in which category-selective signals are the most informative (In load 3 trials, category selectivity is weaker (Daume et al. 2024b)).

The behavioral relevance of this altered neural representation is supported by our mediation analysis, which demonstrated that stimulation-induced changes in FR of category-selective neurons partially mediated the effect of stimulation on RT. This finding indicates that changes in FR account for additional variance in behavior beyond the direct effect of stimulation and suggests that reduced category selectivity of neuronal firing during WM maintenance contributes to impaired behavioral performance. Notably, the indirect effect was smaller than the direct effect. However, this is expected given that the mediation analysis was based on a relatively small number of recorded individual neurons (out of presumably a very large number of persistently active neurons supporting a given WM), a spatially limited stimulation target, and weak stimulation amplitudes. We predict that perturbing a larger neural population, applying stronger or more widespread stimulation, or recording from a greater number of task-relevant neurons would amplify the mediated effect. Additional neural mechanisms, which we did not analyze here, including changes in temporal firing structure, population-level dynamics, or network interactions, are also likely to contribute to the behavioral effects of stimulation.

Notably, the effects of stimulation were specific to the WM maintenance period and were not observed during encoding, when stimulation was absent. This temporal specificity argues against an account based on long-lasting effects of stimulation such as disrupted sensory processing and instead supports the interpretation that hippocampal stimulation interferes with the fidelity of maintained neural representations. Together, these findings indicate that precise category-selective firing patterns, rather than elevated firing alone, are critical for effective WM maintenance. By demonstrating that perturbing the selectivity of persistent neural activity degrades behavior, our results provide evidence in humans that the informational carried by sustained neuronal firing is a key determinant of WM performance.

We note that we deliberately used relatively weak electrical stimulation (1mA, applied through large macro electrodes), a stimulation strength that is weaker than is often used (i.e. see (Parvizi et al. 2013)). This approach was motivated by prior work in primates, which found that localized microstimulation can induce small but reliable behavioral and neuronal changes that can productively be related to each other (Salzman et al. 1992; Celebrini and Newsome 1995). In contrast, we hypothesized (but did not test) that stronger stimulation could cause larger non-specific changes, therefore making it difficult to pin down a specific relationship between the firing of certain neurons and behavior. This approach was successful: despite the weak stimulation, we observed specific effects on category neurons as well as on behavior. Our primary behavioral effect was on RT in load 1, but not accuracy and not load 3. We note that in WM research, RT is commonly used as a sensitive measure of WM quality, with a large body of literature demonstrating that RT slowing is related to weaking of WM representations (Sternberg 1966; Cohen et al. 1997; Luck and Vogel 1997; Donkin and Nosofsky 2012). The absence of an effect on accuracy was likely because subjects optimized accuracy over speed (there was no time pressure). The absence of a significant effect on load 3 we hypothesize is a combination of limited statistical power (the effects are trending in the same direction as load 1) and load 3 recruiting more cognitive resources in other brain areas that can compensate for the disrupted neural activity. Both of these predictions remain to be tested in future work, with our current claims restricted to load 1.

Finally, prior studies of intracranial stimulation in human medial temporal lobe areas, primarily in the context of long-term memory, have shown that stimulation can modulate behavior (Coleshill et al. 2004; Suthana et al. 2012; Fell et al. 2013; Jacobs et al. 2016; Ezzyat et al. 2017; Goyal et al. 2018; Inman et al. 2018; Kucewicz et al. 2018; Jun et al. 2020), but these effects were generally not linked to trial-by-trial single-unit dynamics. Conversely, prior work has demonstrated that electrical stimulation can modulate neuronal firing rates (Cowan et al. 2025), but without directly relating these changes to behavior. By linking stimulation-induced changes in single-neuron activity to WM performance, our study bridges these lines of research and provides a mechanistic framework for understanding how targeted neuromodulation can influence cognition by altering the fidelity of task-relevant neural representations.

### Disruption of Attractor Dynamics

Beyond single-neuron analyses, we show that hippocampal stimulation alters population-level dynamics in a manner consistent with disruption of WM representations in state space. Neural trajectories during WM maintenance normally cluster near stable attractor states, thought to reflect the active representation of the item held in memory (Wang 2001; Wimmer et al. 2014; Kamiński et al. 2017; Panichello et al. 2019; Kamiński and Rutishauser 2020; Penny 2024). In *stimOff* trials, these trajectories remained close to their respective attractors while in *stimOn* trials, they deviated significantly further from them. This increased distance-to-attractor suggests that stimulation perturbs the intrinsic dynamics of the WM network, causing neural representations to become less stable or more ambiguous. Importantly, this effect was specific to hippocampus and VTC, the two regions where selective category neuron activity was also disrupted. Other areas, such as the amygdala, showed intact decoding of stimulus category but no difference in DA values between conditions, supporting a regional specificity of the stimulation effect.

Together, our findings support a theoretical model of WM as an attractor-based system, in which the hippocampus contributes to stabilizing mnemonic representations in neural state space. Electrical stimulation disrupts this stabilization, pushing the network away from optimal configurations and leading to performance declines.

### Implications, Caveats, and Future Directions

Our findings suggest that the mechanisms supporting WM, such as selective persistent activity and attractor dynamics, are distributed across hippocampus and connected cortical areas, such as VTC. Of note, while prominent and strong, our stimulation protocol did not modulate persistent activity in the amygdala. The relevance of amygdala persistent activity for WM therefore remains to be tested, for example by directly stimulating the amygdala.

We note that our claims rely on hippocampal stimulation and we do not know whether stimulation of other parts of the brain would result in similar results. However, given the distributed nature of WM, it seems likely that affected WM-related behavior could be achieved by stimulating various other nodes in the WM network, in particular frontal areas, where stimulation might affect neural activity related to WM control mechanisms. If so, this would further support the notion that WM is supported by a distributed network rather than a single brain area. However, this network appears to be specific: while hippocampal stimulation in our study disrupted persistent activity in some other nodes in the network far away from the stimulation site (hippocampus), it did not disrupt persistent activity in other equally prominent nodes (the amygdala). Future work with different stimulation targets is needed to further dissect the roles of different nodes in the WM network, with the most prominent targets being other brain areas that exhibit persistent activity.

The ways electrical stimulation affects neural activity is complex and non-specific, likely affecting neurons indiscriminately as long as they are close enough to the stimulation electrode. In our recordings only changes in firing rate of neurons with persistent activity mediated effects on RT. Nevertheless, stimulation also modulated the activity of other kinds of neurons and whether such manipulation in turn affected persistently active neurons is not known. A further important open question is to determine whether the extent to which persistent activity disruptions result in impaired WM depends on whether stimulation resulted in off vs. on-manifold disruptions of neural activity (Jazayeri and Afraz 2017).

The present study also demonstrates the feasibility and utility of combining stimulation with single-unit recordings in human subjects. This methodology allows for direct (as opposed to indirect correlational) manipulations that are otherwise difficult to achieve in the human brain and can be used to probe the role of specific neural mechanisms. Future studies could expand this approach to other cognitive domains, such as decision-making or attention, examine how different stimulation parameters (e.g., timing, frequency, amplitude) modulate both behavior and neural codes, and how cell-type specific stimulation can be utilized to precisely target stimulation (Mosher et al. 2020; Lee et al. 2024). Additionally, while we focused here on selective activity in a large time window, future analyses could examine how stimulation affects neural oscillations, spike timing, and interactions between brain regions during WM. This may help further clarify the circuit-level basis of persistent activity and how it supports stable memory representations.

An open question concerns the impact of endogenous disruptions, such as interictal epileptiform discharges, on persistent neuronal activity and behavior. Prior work has shown that such events can modulate single-neuron activity and cognitive performance (Reed et al. 2019), suggesting a potential parallel to the exogenous perturbations examined here. However, assessing these effects is beyond the scope of the present study, representing an important direction for future research.

Finally, our results raise the possibility of therapeutic neuromodulation for disorders of memory. If specific patterns of stimulation can disrupt WM, then appropriately timed or patterned stimulation might be used to enhance or restore it in clinical populations with memory impairments. Understanding how to preserve or augment attractor dynamics may provide a mechanistic basis for such interventions.

## Acknowledgments

We would like to express our deepest gratitude to the patients who volunteered to participate in this study. We thank Clayton Mosher for technical support with the stimulation setup, Nina Long for providing clinical supervision during recording sessions, and the clinical teams at Cedars-Sinai Medical Center (in particular, Jeffrey Chung and Lisa Bateman), Toronto Western Hospital, and Johns Hopkins School of Medicine for patient management and their support of data acquisition. We further thank Tessa Rusch for valuable discussions and Zhongzheng Fu for technical advice. This study was supported by a German National Academy of Sciences Leopoldina Postdoctoral fellowship (to JD), a Center for Neural Science and Medicine at Cedars-Sinai Postdoctoral fellowship (to JD), the BRAIN initiative through the National Institute of Neurological Disorders and Stroke (U01NS103792 and U01NS117839 to UR), and the National Science Foundation (BCS-2219800 to UR).

## Author Contributions

Conceptualization, J.D., J.K. and U.R.; Writing – Original Draft, J.D., and U.R..; Writing – Review & Editing, all authors. Investigation: J.D., M.Y., S.C., C.M.R., I.S., Y.S., J.K., A.C.M.; Formal analysis: J.D. and U.R.; Methodology, J.D., and U.R.; Funding Acquisition, Resources, and Supervision, J.D., U.R. and A.N.M.; Performed surgery, A.N.M., T.A.V., and W.S.A.

## STAR Methods

### Resource availability

#### Lead Contact and Materials Availability

Further information and requests for resources should be directed to the Lead Contact, Ueli Rutishauser.

#### Data and Code Availability

Data will be made available together with example code to reproduce the results on public repositories such as OSF or DANDI upon acceptance, as we commonly do for all our recent publications.

### Experimental Model and Study Participant Details

Thirty patients (32 sessions; 18 females, 11 males, 1 binary; age: 36.8 ± 10.1 years; see **Table S1**) participated in the study. All patients were undergoing intracranial seizure monitoring and surgical evaluation for drug-resistant epilepsy and had Behnke-Fried hybrid electrodes (AdTech Inc.) implanted for clinical purposes (Carlson et al. 2018). Participation in the research was voluntary, and all patients provided informed consent in accordance with the Declaration of Helsinki (Feinsinger et al. 2022). This study was conducted as part of a multi-site NIH BRAIN Initiative consortium involving Cedars-Sinai Medical Center, Toronto Western Hospital, and Johns Hopkins Hospital, and was approved by the Institutional Review Board of each participating institution.

Due to technical constraints, the experimental setup at Johns Hopkins Hospital did not support simultaneous neuronal recordings during electrical stimulation. Consequently, all neural activity analyses were restricted to data collected from 14 patients (15 sessions) at Cedars-Sinai Medical Center and Toronto Western Hospital.

Electrode localization was performed using co-registered pre-operative MRI and post-operative MRI or CT scans, following procedures described previously (Minxha et al. 2020). For visualization purposes, electrode positions were mapped onto the CITI168 atlas brain (Tyszka and Pauli 2016) in MNI152 space (**Fig. 1b**). Apparent electrode placements within white matter or outside target regions reflect the use of a standard template brain for display. Electrodes localized outside of the target regions in native space (N = 4) were excluded from all analyses. Hippocampal sclerosis (see **Table S1**) was assessed by visual inspection of the pre-operative structural MRIs by the attending neurosurgeon operating on a given patient (ratings were 0 = no evidence of sclerosis, 1 = mild, 2 = moderate, 3 = severe sclerosis).

## Method Details

### Task

The task was a modified Sternberg paradigm comprising 140 trials and 280 novel images, as previously used (Daume et al. 2024b, a). Each trial began with a centrally presented fixation cross lasting 0.9–1.2 s (**Fig. 1a**), followed by the encoding phase. Depending on the WM load condition, participants were shown either one image (load 1; 70 trials) or three images in succession (load 3; 70 trials), each displayed for 2 s. After the encoding phase, a maintenance period lasting 2.55–2.85 s followed, during which the word “HOLD” appeared on the screen to signal retention of the encoded items. This was followed by a probe image, which either matched one of the images presented earlier in the same trial (match) or was a non-match image that had been previously seen in earlier trials but not in the current one (see below). Participants were instructed to indicate via button press whether the probe matched one of the images shown during the encoding phase of that specific trial. The probe remained on the screen until a response was made. The response mapping (match vs. non-match) was reversed after half the trials, and this change was communicated to participants during a mid-task break. Responses were recorded using a Cedrus RB-844 response pad (Cedrus Inc.).

All images were novel to each participant and drawn from one of five visual categories: faces, animals, cars (or tools, depending on task version), fruits, and landscapes. Each image appeared at most twice across the entire experiment—once during encoding and once as a potential probe. Importantly, non-match probes were not novel; rather, they were drawn from a pool of images previously seen in earlier trials but from categories not used during the encoding phase of the current trial. This design prevented participants from solving the task based on stimulus novelty alone, thereby enforcing reliance on WM. For patients who participated in multiple sessions, an entirely new set of images was used in each session to ensure that all images remained novel within and across sessions.

### Electrical stimulation

During the WM task, electrical stimulation was delivered to either the left or right hippocampus using biphasic, charge-balanced rectangular pulses. Stimulation was administered at a maximum amplitude of 1.0 mA, a value well below established safety thresholds for short-term intracranial stimulation. To confirm tolerability and avoid triggering afterdischarges, we conducted a pre-experiment ramping procedure in which current was gradually increased from 0.5 mA to 1.0 mA in 0.1 mA increments. In two patients, stimulation amplitude was limited to 0.9 mA or 0.5 mA, respectively, to remain below their individual afterdischarge threshold (see **Table S1**).

Each stimulation trial consisted of 100 biphasic pulses delivered at 50 Hz over a 2 s interval within the WM maintenance period, 0–2000 ms following the offset of the final stimulus. Pulses had a width of 300 μs with a 53 μs inter-phase interval. These parameters have been shown to effectively modulate memory (Suthana et al. 2012; Jacobs et al. 2016). Stimulation was delivered in a bipolar configuration to the two most medial macro-contacts of the Behnke-Fried hybrid electrode implanted in the anterior hippocampus. These contacts were spaced 1.5 mm apart, each with a surface area of 0.059 cm².

The stimulation site (left or right hippocampus) was selected based on clinical diagnosis: stimulation was always applied to the hemisphere with a clinically non-epileptic hippocampus. If both hippocampi were clinically non-epileptic, the hemisphere with more single neuron activity was chosen. Stimulation was administered using the CereStim neurostimulator (Blackrock Neurotech) during a total of 56 trials per session, with trials selected to counterbalance load conditions as well as the visual categories held in WM.

All stimulation sessions occurred at the end of the patient’s hospital stay to ensure a confirmed epilepsy diagnosis and full reinstatement of their antiseizure medication regimen. An experienced neurologist was present during each session and continuously monitored scalp and/or intracranial EEG for evidence of afterdischarges or seizure activity. Patients were blinded to the stimulation condition and did not report perceiving any effects of the stimulation.

## Quantification and Statistical Analysis

### Stimulation artifact rejection and spike sorting

Each hybrid depth electrode included eight microwires from which we recorded broadband local field potential signals ranging from 0.1 to 8,000 Hz, sampled at 32 kHz using the ATLAS recording system (Neuralynx Inc.). All recordings were locally referenced within each recording site by using either one of the eight available micro channels or a dedicated reference channel with lower impedance provided in the bundle. The input range (during the recording or during offline replay of the raw data) was set to 5000 (Cedars-Sinai) or 8000 mV (Toronto Western Hospital).

To minimize the impact of stimulation artifacts on spike sorting quality, we first removed the stimulation pulses from the raw recordings. For this, we processed the broadband data from all recorded microelectrode channels. In each stimulation trial, we identified the precise time points of individual stimulation pulses by detecting their peaks on a channel that remained within the amplifier’s input range (see above). Several control procedures ensured that the number of detected pulses and their inter-pulse intervals matched the expected 50 Hz stimulation pattern (100 pulses with 20 ms in between each pulse), thereby avoiding misclassification.

For each pulse, we interpolated the data from −0.5 ms to +2.0 ms around the pulse peak across all channels simultaneously, effectively removing the stimulation artifact. Visual inspection of all trials and channels confirmed successful artifact removal. Importantly, no data exceeded the amplifier input range following this procedure (see **Fig. 1c** for an example trial). This procedure enabled us to detect spikes in between stimulation pulses.

To maintain comparability between stimulation and non-stimulation conditions and thereby avoid a bias potentially introduced by removing spikes in the stimulation condition only, we applied the exact same cleaning and interpolation procedure to the WM maintenance period in non-stimulated trials as well. To do so, we computed the mean pulse time points (relative to the offset of the last encoded image) across all stimulation trials and interpolated the same −0.5 to +2.0 ms windows around those mean time points in the corresponding segments of non-stimulated trials. Again, visual inspection confirmed the validity of this control procedure.

Following artifact removal, spike detection and sorting were conducted independently on each cleaned and pre-processed microwire using Osort, a semi-automated template-matching algorithm optimized for high-density extracellular recordings (Rutishauser et al. 2006). Prior to spike detection, signals were bandpass filtered between 300 and 3,000 Hz to isolate action potential activity.

To further safeguard against artifact-related contamination, we excluded all detected spikes occurring within an additional 1 ms buffer zone before and after each interpolation window in both stimulation and non-stimulation trials before sorting them into clusters. This ensured that spike sorting was not biased by edge effects from the interpolation. Comprehensive quality metrics for spike sorting, including waveform shape, isolation distance, and cluster separation, are provided in **Fig. S1**. Neurons with a firing rate lower than 0.05 Hz as an average across the entire task were excluded from analysis (19 neurons (2.1%)).

### Selection of neurons

To identify neurons that encoded visual category information, we selected units whose responses following stimulus onset varied systematically across picture categories. For each trial, we quantified neuronal responses by counting spikes within a 200–1,200 ms window after stimulus onset, encompassing all encoding and probe periods. Spike counts were then grouped according to the visual category of the presented stimulus. For each neuron, we performed a one-way ANOVA with picture category as the grouping factor (5 levels), followed by a one-sided, permutation-based t-test comparing the response to the category with the highest mean spike count against all other categories. A neuron was classified as a category neuron if both tests reached statistical significance (p < 0.05, 10,000 permutations; see below). The visual category associated with the maximal firing rate was defined as the neuron’s preferred category.

### Persistent activity of category-selective neurons and interactions with electrical stimulation

To assess whether category neurons exhibited persistent category selectivity during the WM maintenance period and were affected by the electrical stimulation, we conducted analyses of neuronal activity using baseline-normalized FRs (percent change to baseline, using a baseline window of -900 to -300 ms before the first stimulus onset) during the maintenance interval (0–2.5 s after last encoding picture offset). To reduce the bias of outliers, we removed neurons whose FR was higher than 3 SD from the mean across all selected neurons. Areas with less than ten selected neurons for a given analysis were excluded to assure statistical robustness. In all analyses, we employed nested mixed-effects general linear models (GLMs) with neuronID nested into patientID to robustly model the hierarchical nature of the recorded data, accounting for repeated measures and subject-level variability (Aarts et al. 2014).

To assess persistent category selectivity during the WM maintenance, we analyzed all trials in which patients responded correctly. We employed nested mixed-effects GLMs with *preference* (2 levels: preferred, non-preferred; categorical) as a fixed effect, and random intercepts for neuronID nested within patientID. The model specification was:

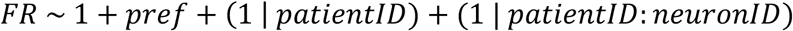

To investigate whether electrical stimulation modulated this persistent activity, we extended the GLM to include *stimulation* (2 levels: stimOn, stimOff; categorical) and the interaction between *preference* and *stimulation*, using the same nested design and restricting analysis to correct trials only:

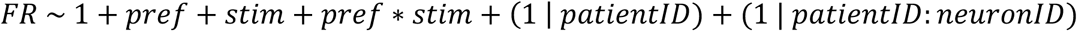

The stimulation x preference interaction term provided a formal statistical test of whether category selectivity differed between stimulation ON and OFF conditions. To confirm that any observed interaction effects were specific to the WM maintenance period and not present during stimulus encoding, i.e., when no stimulation occurred, we performed a control analysis using the same model structure but based on FRs from the window used to define category selectivity (0.2–1.2 s after onset of the first image).

### Mediation analysis

We tested whether the effect of electrical stimulation on behavior (**Fig. 1**) was mediated by stimulation-induced changes in the FR of category-selective neurons, which would be consistent with a direct contribution of neuronal firing to WM–related behavioral changes. To this end, we performed a mediation analysis (MacKinnon et al. 2007; Imai et al. 2010) using the *mediation* package in R (Tingley et al. 2014). The mediation analysis decomposes the total effect of stimulation on RT into an indirect effect mediated by neuronal FR and a direct effect not explained by FR. Specifically, we estimated the average causal mediation effect (ACME), corresponding to the effect of stimulation on RTs transmitted via changes in FR, and the average direct effect (ADE), corresponding to the effect of stimulation on RTs not mediated by FR. The total effect was defined as the sum of ACME and ADE.

Based on the behavioral results and the observed stimulation-related changes in FR, we restricted this analysis to load 1 trials, which exhibited the strongest stimulation effects on RTs. Furthermore, only trials in which non-preferred images were maintained were included, as significant FR modulation by stimulation was observed most strongly under this condition. Because the mediation analysis relied solely on non-preferred trials, we further excluded neurons that did not exhibit a clearly defined single preferred category (13 neurons; see **Fig. S2e** for an example). This exclusion ensured that the non-preferred condition did not include stimuli eliciting some residual selectivity for a given neuron, which could otherwise confound estimates of FR modulation in the non-preferred condition.

The mediation analysis was implemented using two linear mixed-effects models. First, we modeled the effect of stimulation on neuronal firing rate across the selected neurons:

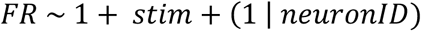

Second, we modeled the effects of FR and stimulation on behavioral response times:

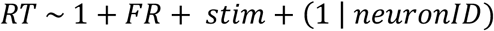

Statistical inference on ACME, ADE, and the total effect was performed using the mediate function of the *mediation* package in R, specifying *stimulation* as the treatment variable and *FR* as the mediator. Confidence intervals and p-values were obtained via 2,000 nonparametric bootstrap simulations.

As with all mediation analyses, estimation of ACME and ADE relies on the assumption of no unmeasured confounders of the mediator–outcome relationship. While this assumption cannot be directly tested, it is partially mitigated in the present study by the experimental manipulation of stimulation and the restriction to a homogeneous trial condition.

### Population decoding

To evaluate the effect of electrical stimulation on neural activity across brain regions, we used linear support vector machine (SVM) classifiers to decode *stimOn* versus *stimOff* trials based on spike counts during the WM maintenance period. This analysis was conducted separately for each recorded region using correct trials only.

For each brain area, we constructed a pseudo-population by pooling neurons across all sessions. Spike counts during the maintenance period served as input features for the decoder. Trial counts were balanced across conditions by randomly subsampling trials in each session to match the minimum number of available trials. We performed 10-fold cross-validation to assess decoding accuracy, normalizing spike counts for each neuron across trials within the training set in each fold to prevent data leakage. To account for variability due to trial selection, the subsampling procedure was repeated 50 times. Final decoding accuracy was computed as the average across repetitions, and error bars in **Fig. 1h** represent the standard deviation of accuracy across these 50 subsampled repetitions.

To assess statistical significance and construct a null distribution, we performed a label-shuffling control. Trial labels (*stimOn* vs. *stimOff*) were randomly permuted, and the same decoding procedure (including subsampling and cross-validation) was repeated 500 times for each brain area. This yielded an empirical distribution of decoding accuracies under the null hypothesis of no relationship between neural activity and stimulation condition.

### State-space attractor analysis

Previous work has shown that during WM maintenance, neural population activity tends to reside near low-dimensional attractor states in neural state space, and that proximity to these attractors correlates with WM performance on a trial-by-trial basis (Kaminski et al., 2017). To investigate how electrical stimulation affected population dynamics in state space—specifically the distance to attractor states—we applied demixed principal component analysis (dPCA) (Kobak et al. 2016), using picture category as the marginalized variable.

FRs from all recorded neurons were binned using 200 ms wide sliding windows with 1 ms steps, allowing for fine-grained temporal resolution. We then constructed a pseudo-population by pooling neurons across all recording sessions. Given that stimulation effects on neural activity were similar in hippocampus and VTC, neurons from both areas were combined for this analysis. The demixing axes (basis functions) were computed using FRs from a window 200–1,200 ms following the onset of the first image in the encoding phase (encoding 1). This ensured that the WM maintenance period remained independent and unbiased for downstream analyses. To eliminate potential ambiguity due to multi-item interference, only correct load 1 trials were included. To prevent overfitting during dimensionality reduction, we used regularization and selected the optimal lambda parameter via cross-validation (Kobak et al. 2016; Kamiński et al. 2017).

We rank-ordered the demixed principal components (dPCs) by their explained variance and selected the four components (1, 2, 3, and 5) that accounted for the most variance attributable to category identity, consistent with previous studies (Kamiński et al. 2017). The resulting low-dimensional space allowed us to trace population-level trajectories during WM maintenance. To quantify proximity to attractor states, we first defined a set of attractors *A_k_*, one for each of the five visual categories maintained in WM. Each attractor *A_k_* was defined as the time-averaged neural state observed during the maintenance period for *stimOff* trials in which category k was remembered.

The distance-to-attractor (DA) metric was computed for each trial based on the relative distance of the neural trajectory to its corresponding attractor, normalized by its distances to all other attractors (Kamiński et al. 2017). Specifically, for each time point t during the maintenance period:

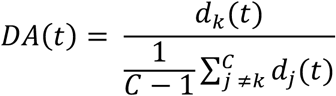

Where *d_j_(t) = ||p(t) – A_j_||* is the Euclidean distance between the neural trajectory *p(t)* at time *t* and attractor *A_j_*, and C is the number of attractors (*C = 5*). DA values were computed at each time point and then averaged across the maintenance period to obtain a single scalar DA value per trial. A DA value less than 1 indicates that the neural trajectory was closer to its correct attractor than to the average distance from all others, reflecting strong category-specific encoding in state space.

### Statistics

Throughout the manuscript, statistical comparisons between conditions were assessed using t-tests, ANOVAs, or nested mixed-effects GLMs. GLMs were implemented using fitglme.m in MATLAB. For t-tests and ANOVAs, we employed non-parametric permutation statistics using the statcond.m function from the EEGLAB toolbox. This approach does not assume specific underlying data distributions and was conducted with 10,000 permutations unless stated otherwise. Reported *t* and *F* values, which are based on parametric assumptions, are included for reference only. Unless otherwise noted, error bars in all figures represent the standard error of the mean (s.e.m.).

