## Supplementary information for "Disrupting selective persistent activity with electrical stimulation impairs human working memory"

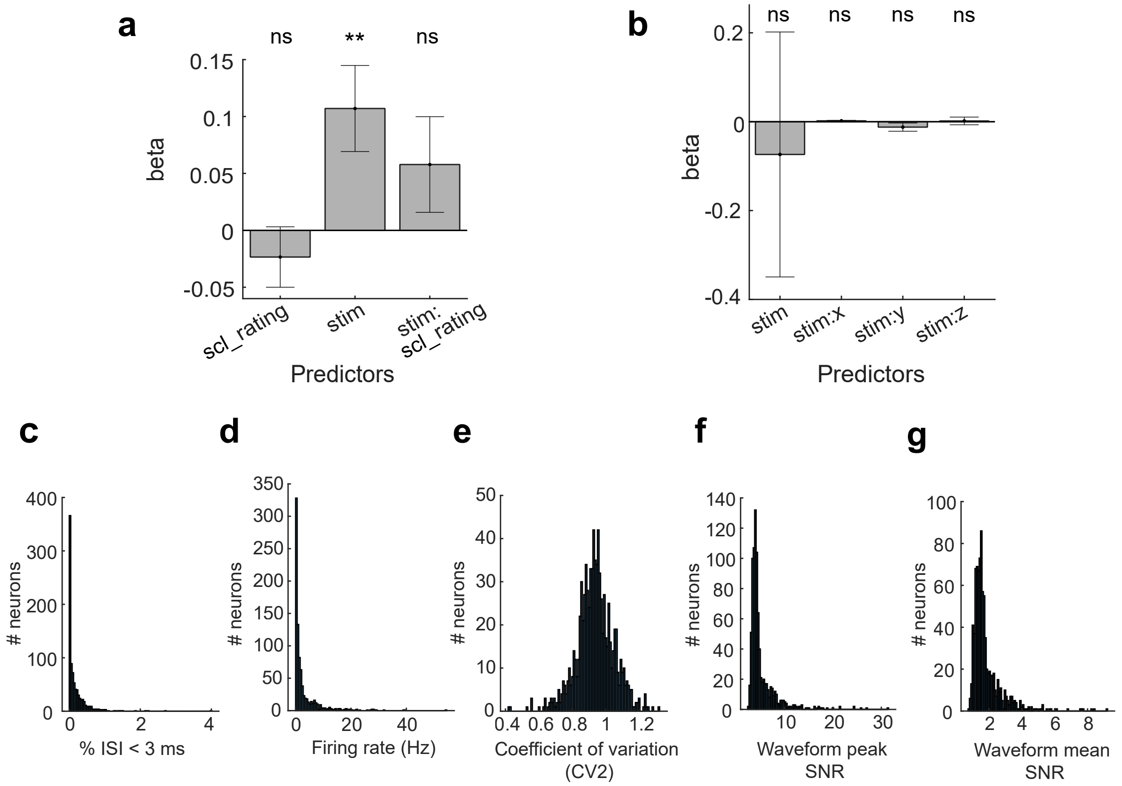


**Figure S1. Behavior of patients included in neuronal analysis and spike-sorting quality metrics for all identified putative single units. (a)** A mixed-effects GLM with factors sclerosis rating and stimulation on/off revealed no significant interaction between stimulation and hippocampal sclerosis rating (see Table S1) on reaction times, suggesting that the severity of sclerosis did not account for the observed stimulation-related changes in behavior. **(b)** Similarly, anatomical location of stimulation along the x-, y-, or z-axis in MNI space did not modulate the magnitude of the stimulation effect on reaction times as indicated by no significant interaction with stimulation on/off. **(c)** Proportion of inter-spike intervals (ISI) below 3 ms. **(d)** Average firing rate. **(e)** Coefficient-of-variation. **(f)** Signal-to-noise ratio (SNR) for the peak of the mean waveform across all spikes as compared to the standard deviation of the background noise. **(g)** Mean SNR of the waveform.

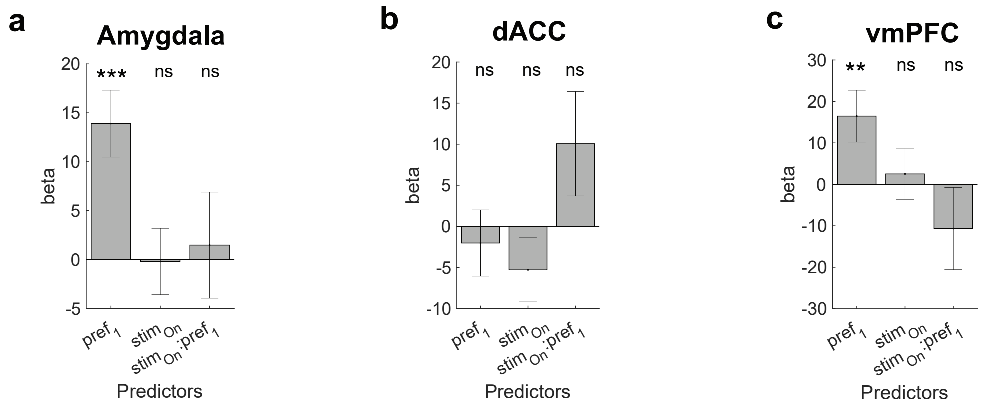


d

| | Estimate | 95% CI Lower | 95% CI Upper | p-value |
|---|---|---|---|---|
| ACME | -9.17e-05 | -5.06e-04 | 2.86e-04 | 0.64 |
| ADE | 3.92e-02 | 2.98e-02 | 4.79e-02 | < 2e-16 |
| Total Effect | 3.91e-02 | 2.96e-02 | 4.78e-02 | < 2e-16 |
| Proportion Mediated | -1.98e-03 | -1.36e-02 | 7.63e-03 | 0.64 |

Mediation results

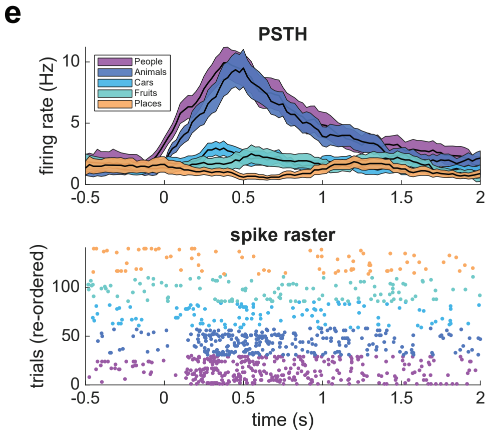


**Figure S2. Effect of electrical stimulation on category neuron activity across additional brain regions.** Mixed-effects GLMs revealed no significant interaction between *stimulation* and *preference* in **(a)** amygdala, **(b)** dACC, and **(c)** vmPFC. Pre-SMA was excluded from this analysis due to an insufficient number of identified category neurons for statistically robust inference. **(d)** Results of the mediation analysis including all non-selective neurons in the hippocampus and VTC. In contrast to selective neurons, we did not observe a significant indirect effect (ACME) of FR using this group of neurons. **(e)** An example neuron from the VTC that showed selectivity to two different picture categories. ***p < 0.001, **p < 0.01; ns = not significant; mixed-effects GLMs.

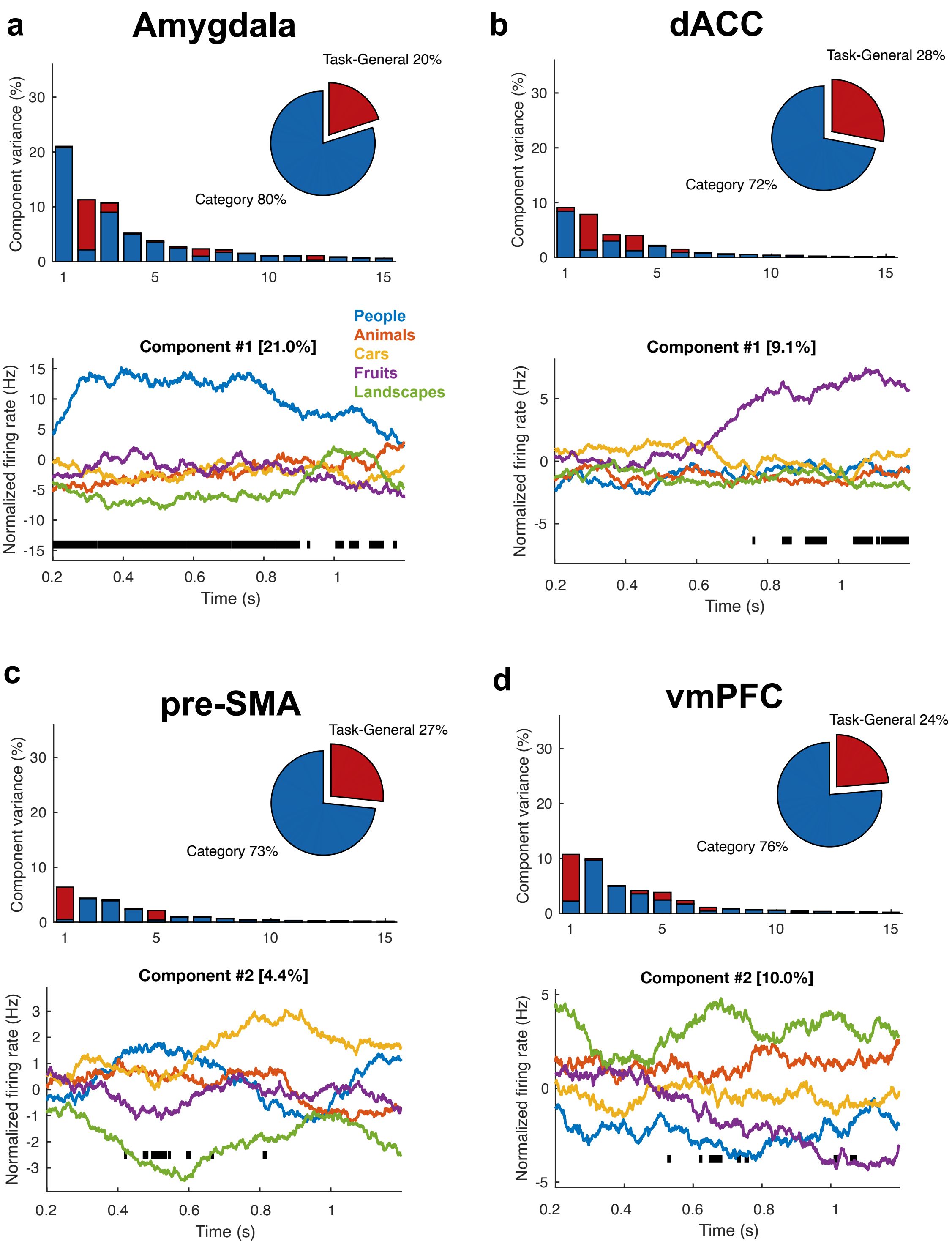


**Figure S3. Population-level category decoding in additional brain regions.** dPCA revealed robust above-chance separation of category information in **(a)** amygdala, but not in **(b)** dACC, **(c)** pre-SMA, or **(d)** vmPFC. Line plots show firing rates projected onto the most informative category-selective components.

| Session ID | sex | age | SOZ | Stim area | MTS | Stim amp (mA) | Hippo | Amy | VTC | preSMA | dACC | vmPFC |
|---|---|---|---|---|---|---|---|---|---|---|---|---|
| **P67cs** | F | 38 | Bilateral mesial temporal | LH | 3 | 1.0 | 2 | 12 | 0 | 0 | 5 | 25 |
| **P69cs** | F | 41 | Not localized, broad onset | LH | 0 | 1.0 | 14 | 10 | 0 | 0 | 17 | 12 |
| **P76cs** | F | 24 | Not localized | RH | 0 | 1.0 | 3 | 9 | 5 | 6 | 8 | 1 |
| **P78cs** | F | 54 | Right anterior temporal | RH | 0 | 1.0 | 6 | 18 | 5 | 0 | 0 | 0 |
| **P79cs** | F | 42 | Right anterior lateral temporal neocortical | RH | 1 | 1.0 | 15 | 29 | 29 | 10 | 15 | 21 |
| **P80cs** | O | 24 | Not localized | RH | 1 | 1.0 | 17 | 41 | 5 | 12 | 10 | 3 |
| **P81cs** | F | 28 | Left mesial temporal lobe | RH | 0 | 1.0 | 3 | 10 | 27 | 9 | 4 | 8 |
| **P82cs** | M | 42 | bitemporal | RH | 0 | 0.9 | 16 | 8 | 2 | 1 | 4 | 0 |
| **P86cs** | F | 39 | Bifrontal: no definite localization | LH | 0 | 1.0 | 7 | 10 | 9 | 20 | 18 | 8 |
| **P90cs** | M | 32 | Left mesial hipp | RH | 0 | 1.0 | 14 | 20 | 5 | 5 | 17 | 15 |
| **P106cs** | M | 37 | Left hipp plus neocortical (multifocal) | RH | 0 | 1.0 | 6 | 13 | 1 | 18 | 6 | 18 |
| **P107cs** | F | 65 | Left mesial temporal | RH | 0 | 1.0 | 14 | 20 | 30 | 35 | 45 | 17 |
| **TWH232** | M | 30 | Left mid temporal | LH | 0 | 1.0 | 3 | 0 | 0 | 0 | 0 | 0 |
| **TWH232_2** | M | 30 | Left mid temporal | LH | 0 | 1.0 | 3 | 0 | 0 | 0 | 0 | 0 |
| **TWH233** | F | 25 | Left middle insula | RH | 0 | 1.0 | 18 | 21 | 0 | 0 | 0 | 0 |
| **JH_PY23N003** | F | 27 | bilateral temporal region | LH | 1 | 1.0 | 0 | 0 | 0 | 0 | 0 | 0 |

| | | | | | | | | | | | | |
|---|---|---|---|---|---|---|---|---|---|---|---|---|
| **JH_PY23N007** | M | 41 | Right temporo-parieto-occipital region | RH | 0 | 1.0 | 0 | 0 | 0 | 0 | 0 | 0 |
| **JH_PY23N009** | F | 57 | Right amygdala and hippocampus, | LH | 0 | 1.0 | 0 | 0 | 0 | 0 | 0 | 0 |
| **JH_PY23N011** | M | 40 | right fronto-temporal | RH | 1 | 1.0 | 0 | 0 | 0 | 0 | 0 | 0 |
| **JH_PY23N017** | F | 44 | right anterior temporal lobe | RH | 0 | 1.0 | 0 | 0 | 0 | 0 | 0 | 0 |
| **JH_PY23N019** | F | 42 | Primary motor and primary sensory cortex | LH | 0 | 1.0 | 0 | 0 | 0 | 0 | 0 | 0 |
| **JH_PY23N022** | F | 31 | right temporal lobe region | RH | 2 | 1.0 | 0 | 0 | 0 | 0 | 0 | 0 |
| **JH_PY23N023** | F | 44 | right anterior temporal lobe | RH | 2 | 1.0 | 0 | 0 | 0 | 0 | 0 | 0 |
| **JH_PY23N027** | F | 36 | left amygdala | LH | 1 | 1.0 | 0 | 0 | 0 | 0 | 0 | 0 |
| **JH_PY24N002** | M | 36 | left temporal lobe | LH | 0 | 1.0 | 0 | 0 | 0 | 0 | 0 | 0 |
| **JH_PY24N004** | F | 24 | right temporal lobe | RH | 1 | 1.0 | 0 | 0 | 0 | 0 | 0 | 0 |
| **JH_PY24N005** | M | 32 | left anterior thalamic nuclei | LH | 1 | 1.0 | 0 | 0 | 0 | 0 | 0 | 0 |
| **JH_PY24N010** | M | 39 | right temporal lobe | RH | 0 | 1.0 | 0 | 0 | 0 | 0 | 0 | 0 |
| **JH_PY24N012** | M | 34 | left temporal pole | LH | 0 | 1.0 | 0 | 0 | 0 | 0 | 0 | 0 |
| **JH_PY24N013** | F | 21 | right mesial temporal structures | LH | 0 | 1.0 | 0 | 0 | 0 | 0 | 0 | 0 |
| **JH_PY25N001** | M | 34 | Left temporal lobe | LH | 0 | 0.5 | 0 | 0 | 0 | 0 | 0 | 0 |
| **JH_PY25N001_2** | M | 34 | Left temporal lobe | LH | 0 | 0.5 | 0 | 0 | 0 | 0 | 0 | 0 |

**Table S1. Patient demographics and neuron count per area.** M = male; F = female; O = binary. L/RH = left/right hippocampus. MTS = Mesial Temporal Sclerosis Rating (of the stimulated hippocampus; 0 = no evidence of sclerosis, 1 = mild, 2 = moderate, 3 = severe sclerosis).
